# The Swiss Primary Ciliary Dyskinesia registry: objectives, methods and first results

**DOI:** 10.1101/450700

**Authors:** Goutaki Myrofora, Eich Marc, Florian S. Halbeisen, Barben Juerg, Casaulta Carmen, Clarenbach Christian, Gaudenz Hafen, Latzin Philipp, Regamey Nicolas, Lazor Romain, Tschanz Stefan, Zanolari Maura, Maurer Elisabeth, Kuehni E. Claudia, Swiss PCD Registry (CH-PCD) Working Group

## Abstract

Primary Ciliary Dyskinesia (PCD) is a rare hereditary, multi-organ disease caused by defects in ciliary structure and function. It results in a wide range of clinical manifestations, most commonly in the upper and lower airways. Central data collection in national and international registries is essential to studying the epidemiology of rare diseases and filling in gaps in knowledge of diseases such as PCD. For this reason, the Swiss Primary Ciliary Dyskinesia Registry (CH-PCD) was founded in 2013 as a collaborative project between epidemiologists and adult and paediatric pulmonologists.

The registry records patients of any age, suffering from PCD, who are treated and resident in Switzerland. It collects information from patients identified through physicians, diagnostic facilities, and patient organisations. The registry dataset contains data on diagnostic evaluations, lung function, microbiology and imaging, symptoms, treatments, and hospitalizations.

By May 2018, CH-PCD has contacted 566 physicians of different specialties and identified 134 patients with PCD. At present this number represents an overall 1 in 63,000 prevalence of people diagnosed with PCD in Switzerland. Prevalence differs by age and region; it is highest in children and adults younger than 30 years, and in Espace Mittelland. The median age of patients in the registry is 25 years (range 5-73), and 49 patients have a definite PCD diagnosis based on recent international guidelines. Data from CH-PCD are contributed to international collaborative studies and the registry facilitates patient identification for nested studies.

CH-PCD has proven to be a valuable research tool that already has highlighted weaknesses in PCD clinical practice in Switzerland. Development of centralised diagnostic and management centres and adherence to international guidelines are needed to improve diagnosis and management—particularly for adult PCD patients.

## Introduction

Primary Ciliary Dyskinesia (PCD) is a rare hereditary, multi-organ disease. Genetic mutations cause defects in ciliary structure and function that result in a wide range of clinical manifestations [1]. In the upper and lower airways, impaired mucociliary clearance leads to recurrent and chronic infections such as rhinosinusitis, otitis, chronic wet cough and pneumonia, and subsequently to more severe manifestations in many patients that include irreversible lung damage, bronchiectasis, and hearing impairment [2-5]. Cilia play an important role in organ placement in utero and about half of PCD patients present with situs inversus, while 10% have other laterality defects (situs ambiguous). [6] More rarely, about 5% of patients have simple or complex cardiovascular malformations. [7] Other organ systems that may be affected in patients with PCD include the reproductive system, with many male patients reporting infertility because of immotile or dysfunctional sperm. In women, the fallopian tubes are lined with ciliated cells that play a role in the transport of gametes and embryos; thus some female patients with PCD report fertility problems such as ectopic pregnancies. Other organs can be affected, and hydrocephalus, retinitis pigmentosa, and renal cysts have been occasionally reported.

Research in PCD, as in other rare diseases, faces challenges. Currently, 39 genes have been found to be associated with the disease in 65-70% of patients.[8] Yet in spite of increasing understanding of its genetic basis and knowledge about PCD, the disease has no dedicated International Classification of Diseases revision 10 (ICD-10) code. Otherwise routine epidemiological data such as PCD mortality and hospitalisations are therefore unavailable. Additionally, PCD is difficult to diagnose. Lacking a single, standard diagnostic test,[9] diagnosis of PCD requires a combination of tests, some of which require high expertise, that are based on evidence-based guidelines [9,10].

Despite advances in PCD diagnostics, many patients are still misdiagnosed or diagnosed late in life so prevalence is not accurately known. Prevalence of PCD is estimated to be around 1 in 10,000, but the figure varies considerably between studies.[1] This is an unavoidable product of low numbers. Even large hospitals care for few PCD patients, and single-centre studies have small sample sizes. Care is decentralized and differs between countries—or even within the same country. In Switzerland, we found that most clinics care for fewer than three PCD patients [11].

Centralised collection of data in national and international registries is essential to raising patient numbers in the epidemiological study of rare diseases such as PCD, and filling in gaps in knowledge. Centralised data collection allows comparison and standardisation of care and study of the long-term prognosis and quality of life of PCD patients, and thus was CH-PCD founded in 2013 as a collaborative project between epidemiologists and adult and paediatric pulmonologists. Beginning as a pilot project in the canton of Bern, in 2014 CH-PCD was extended to include all regions in Switzerland. The data centre of CH-PCD is located at the Institute of Social and Preventive Medicine (ISPM) at the University of Bern and contributes data to the international PCD registry [12] and other international studies such as the international PCD (iPCD) cohort [13].

Here we describe the objectives and methodology of the CH-PCD, present initial results, and give an overview of current and ongoing projects.

## Objectives of the Swiss PCD Registry

CH-PCD collects information on diagnosis, symptoms, treatment, and follow-up of patients with PCD in Switzerland, and provides data for national and international monitoring and research. In particular, it aims to

- Identify all patients diagnosed with PCD in Switzerland
- Collect population-based data: incidence, prevalence, time and regional trends
- Document diagnostic evaluations, treatments, and participation in clinical trials
- Document the clinical course of PCD, quality of life, morbidity, and mortality
- Establish a research platform for clinical, epidemiological, and basic research

## Materials and methods

### Study design

CH-PCD is a patient registry (clinicaltrials.gov registration number: NCT03606200). At baseline, when a patient is included in the registry, it retrospectively collects all available data since birth and then follows patients until death or loss to follow up, collecting prospective data at regular intervals.

### Study population

CH-PCD registers Swiss residents of any age who suffer from PCD. Since PCD diagnostics have evolved through the years, CH-PCD includes patients diagnosed in different ways. Some patients have been diagnosed with a combination of tests in accordance with current recommendations [10,14], while diagnosis of others can be based solely on strong clinical suspicion that will need to be confirmed when the diagnostic algorithm is completed or when improved diagnostic methods become available.

### Patient identification procedures

The CH-PCD collects information from patients identified by physicians, diagnostic facilities, patient organisations such as the Kartagener Syndrom und Primäre Ciliäre Dyskinesie e.V. (http://www.kartagener-syndrom.org/), and the Swiss group for Interstitial and Orphan Lung diseases (the SIOLD Registry, http://www.siold.ch). The CH-PCD team identifies physicians working in specialties related to PCD management—particularly paediatric and adult pulmonologists, otolaryngologists (ENT specialists), and fertility specialists—in hospitals, clinics, and private practices. We contact them by email, send them the study information, and ask whether they treat any PCD patients (Figure 1). Those who do not reply receive phone calls. Physicians who report PCD patients receive a baseline reporting form, which enquires about the information necessary to identify and contact the patient (name, address, date of birth, date and type of diagnosis, and last date of follow-up). Patients thus identified are contacted by their physician or, per the physician’s request, by the CH-PCD team directly. In the first case, the physician provides and explains the study information and an informed consent form, at the next, follow-up appointment. When the ISPM team contacts the patient directly, these documents are sent by post. Study information and consent forms are available in three languages (French, German, and Italian).

**Figure 1.**
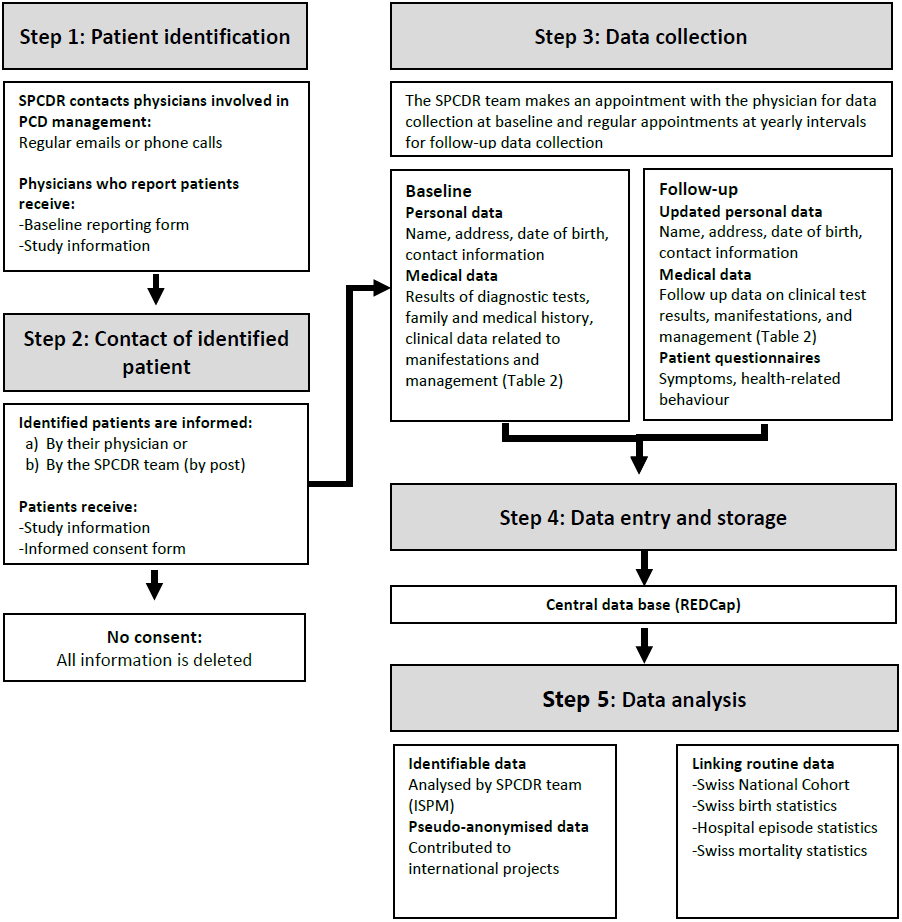
Schematic chart of patient identification and data collection for the Swiss PCD registry

### Data sources

Detailed data are collected from patients who give informed consent or assent, upon reception of the patient registration form (Figure 1). The CH-PCD team makes an appointment with the physician for data collection. At the time of inclusion in the registry, the following documents are thoroughly searched for any retrospectively available relevant information: referral letters, discharge letters, paper and electronic medical charts, lung function measurements, laboratory results, imaging and diagnostic tests, drug prescriptions, physiotherapy prescriptions, and surgery reports. After inclusion in the registry, further information from routine follow-up visits and data related to emergency visits and hospitalizations are collected prospectively from the same sources at yearly intervals.

### What data is collected

CH-PCD collects information on demographic characteristics such as age and sex, diagnostic tests, and clinical data about manifestations and management of the disease. It regularly collects follow-up data on growth, lung function, clinical manifestations from all affected organ systems, microbiology, imaging, lab results, therapeutic interventions including surgery and physiotherapy, and hospitalisations. It also collects information on neonatal symptoms related to the disease and on the symptoms that led to referral and PCD diagnosis. Table 1 contains a detailed overview of the data CH-PCD collects.

**Table 1.** Description of data collected in the Swiss PCD registry

|  | Type of data | Collected variables | Data collection |
| --- | --- | --- | --- |
| 1) | General information | patient demographic data date of birth, sex, ethnicity, date of last follow up, date last known alive) and information related to registration and treating physicians (date of informed consent, current and past treating physicians) | cross-sectional |
| 2) | Diagnostics | diagnostic status of PCD, date of diagnosis, dates, and results of diagnostics test performed (nNO testing, VM, EM, genetic testing) and dates and results of tests performed to exclude other diagnoses (cystic fibrosis tests) | longitudinal |
| 4) | Clinical manifestation leading to diagnosis | information on past symptoms and signs from the upper and lower airways (cough, upper and lower respiratory infections, hearing impairment) and other systems (laterality defects, congenital heart defects, brain cilia dysfunction, retinitis pigmentosa, renal problems, fertility problems) from the referral letters for diagnostic evaluation | cross-sectional |
| 5) | Neonatal period | birth somatometric measurements, symptoms and signs of neonatal period (neonatal respiratory distress syndrome, neonatal cough, neonatal rhinitis), need for intensive care and ventilation | cross-sectional |
| 6) | Growth and lung function | lung function date, somatometric measurements, spirometric measurements, other lung function tests (body plethysmography, breath washout) | longitudinal |
| 7) | Clinical manifestations | date of clinical visit, information on upper and lower respiratory symptoms and signs (cough, upper and lower respiratory infections, hearing impairment) and symptoms and signs from other affected systems (laterality defects, congenital heart defects, fertility problems) and active or passive smoking during the follow-up period since last visit | longitudinal |
| 8) | Therapy | date of prescription, information on inhaled and nasal medication (bronchodilators, inhaled corticosteroids, nasal sprays), antibiotics prescription, oxygen therapy, mechanical ventilation, physiotherapy and vaccinations | longitudinal |
| 9) | Hospitalisations and surgical interventions | information on hospitalisation (clinic, reason for hospitalisation, diagnosis, date of discharge), date and type of surgery | longitudinal |
| 10) | Microbiology | date of microbiology examination, type of sample, and isolated bacteria | longitudinal |
| 11) | Imaging | date and findings of chest and sinus imaging (X-Ray, CT, and MRI), information on location of the findings and information on other types of imaging | longitudinal |
| 12) | Other tests | date and results of other performed tests (blood gas tests, atopy tests) | longitudinal |
| 13) | Family history | number of siblings, PCD affected siblings and other family members, patient ID of PCD affected family members, and relationship of affected family members | cross-sectional |

nNO, nasal nitric oxide; VM, videomicroscopy; EM, electron microscopy; CT, computed tomography; MRI, magnetic resonance imaging

### Ethical approval/ Data protection

In 2013, CH-PCD obtained special registry permission from the Expert Committee for Professional Secrecy in Medical Research of the Federal Office of Public Health (FOPH). After the new Human Research Act came into effect in Switzerland, CH-PCD obtained an authorisation from the Cantonal Ethics Committee of Bern in 2015 (KEK-BE: 060/2015). The authorisation is valid for data collection all over Switzerland and includes the retrospective registration of data from deceased patients and patients lost to follow-up. For patients lost to follow-up, the registry collects basic data and last known address. Addresses are updated in a uniform way by contacting community registration offices, and as soon as current addresses of patients are found patients are contacted, informed, and asked for consent.

Patient data are collected at hospitals, clinics, and in private practices, and any additional patient information is handled via post or a secure e-mail address.Data collection is performed by the CH-PCD team using a local copy of the database stored on an external hard drive with strong encryption (AES 256). In the event photocopies of data are made, they are destroyed after the data have been entered into the CH-PCD database. Personal information (names and addresses) is kept strictly separate from the clinical information of the CH-PCD database and is stored in a separate database.

The web-based clinical database uses Research Electronic Data Capture (REDCap) developed at Vanderbilt University (Nashville, TN, USA) [15]. REDCap supports all necessary security requirements, is widely used in the academic research community, and allows data extraction in various formats as well as linkage with international PCD research projects such as the iPCD cohort. Access to the ISPM server and the CH-PCD database is restricted to selected persons working on the registry. Data in paper form are securely locked and can only accessed by the CH-PCD team. Daily, weekly, and monthly back-ups are performed and securely stored on the ISPM servers.

### Data quality and analyses

CH-PCD data from each patient are manually checked for completeness, plausibility, and consistency. Ambiguities are resolved in collaboration with treating physicians at the time of data collection or after the manual quality check. Data extractions for embedded national or international studies are routinely checked for quality assurance. Basic data are presented in the annual CH-PCD report and sent to all collaborating physicians and patient organizations. For each planned analysis, eligible data are validated and anonymized. For national analyses, CH-PCD data are analysed at ISPM Bern. Data are also contributed to international collaborative analyses. Since not all patients have diagnostic information that is up to current standards, all analyses are stratified by level of diagnostic certainty. Appropriate statistical methods are selected based on the research question and all statistical analyses are performed using STATA and R.

### Funding

The setting up of CH-PCD (salaries, consumables, equipment) was funded by several Swiss funding bodies that include the Lung Leagues of Bern, St. Gallen, Vaud, Ticino, and Valais, and the Kantonalbernischer Hilfsbund. PCD research at ISPM Bern is funded by the Swiss National Science foundation (SNF 320030_173044). CH-PCD also participates in the EU funded BEAT-PCD COST Action (BM1407) [16,17].

## Results

### Patient identification

A special effort has been made to contact all paediatric and adult pulmonologists, otolaryngologists, and fertility specialists in the cantons of Appenzell Ausserrhoden, Appenzell Innerrhoden, Basel-Landschaft, Basel-Stadt, Bern, Fribourg, Glarus, Graubünden, Jura, Neuchâtel, Schaffhausen, St.Gallen, Thurgau, Ticino, Valais, Vaud, and Zürich. An additional effort to contact physicians from other specialties such as cardiology and ophthalmology, who may treat rarer extrapulmonary manifestations of PCD patients, as well specialists in neonatology and radiology who may have been involved in management of patients began in the canton of Bern in spring 2018. Among the 566 physicians contacted by May 2018, 418 (74%) responded to confirm whether they had or had not managed patients with PCD. The response rate was higher among adult and paediatric pulmonologists and ENT specialists, who are more involved in disease management (Table 2). The physician respondents identified 134 patients who were registered in CH-PCD. Three patients refused consent and were not included in the registry. Detailed data have already been collected for 78 patients. For the remaining 53 patients, basic demographic and diagnostic data are currently available. The numbers of patients identified by canton varied from zero to 46, and most physicians reported either no or at most five patients. Patient identification of retrospectively diagnosed or newly diagnosed patients is ongoing.

**TABLE 2.** Patient identification for the SPCDR: contacted physicians, response rate, and identified patients by medical specialty

| **Specialty** | **Contacted physicians** | **Responding physicians** | **Response rate** | **Identified patients** |
| --- | --- | --- | --- | --- |
| Pulmonologists | 152 | 120 | 79% | 54 |
| Paediatric pulmonologists | 57 | 44 | 77% | 75 |
| ENT specialists | 164 | 132 | 80% | 3 |
| Fertility specialists/urologists | 67 | 41 | 61% | 0 |
| Cardiologists | 37 | 23 | 62% | 0 |
| Ophthalmologists | 37 | 17 | 46% | 0 |
| Neurologists | 19 | 16 | 84% | 0 |
| Nephrologists | 6 | 6 | 100% | 0 |
| Radiologists | 15 | 9 | 60% | 0 |
| General paediatricians/<br>general practitioners* | 12 | 10 | 100% | 2 |
| **Total** | **566** | **418** | **74%** | **134** |

\*General paediatricians and general practitioners were not contacted systematically, but only in selected patients identified through the *Kartagener Syndrom und Primäre Ciliäre Dyskinesie e.V.* patient organization or when prompted by other physicians

### Prevalence of diagnosed disease

With an assumed prevalence of 1 to 10,000 [1], 842 PCD patients are expected to live in Switzerland (population of 8,419,550; BSF 2016). The 134 patients identified to date result in a prevalence of 1 in 63,000 (which represents about 1 in 6 of the expected number of Swiss PCD patients). This prevalence varied by age (Figure 2). We found a much higher prevalence of known cases in children aged 0-10 (1 in 44,000), adolescents aged 11-20 (1 in 24,000), and young adults (1 in 36,000) compared to adults older than 30 (1 in 104,000– 130,000). Prevalence also differed by geographic region (Figure 3), with most cases treated in Espace Mittelland—the cantons of Bern, Fribourg, Jura, Neuchâtel, and Solothurn (prevalence 1 in 37,000), and the fewest in Ticino (prevalence 1 in 177,000).

**Figure 2.**
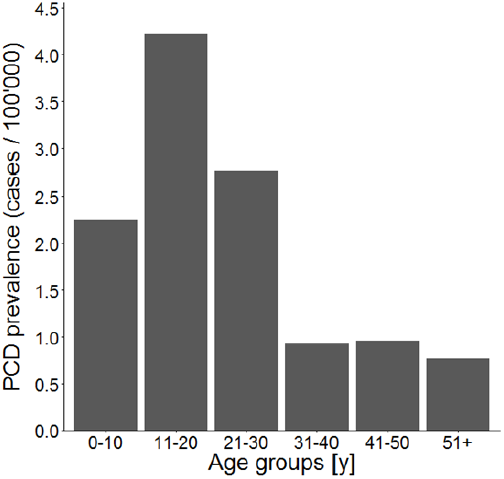
Prevalence of PCD diagnosed in Switzerland by age group Calculated by dividing the number of patients identified in the Swiss PCD registry by the number of the people in the respective age group living in Switzerland. Prevalence is presented as cases/100,000 inhabitants.

**Figure 3.**
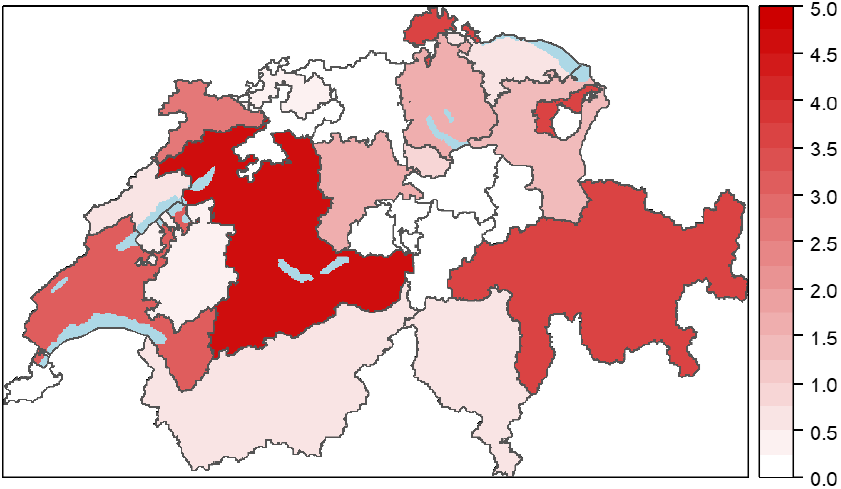
Prevalence of diagnosed PCD patients treated in the different cantons of Switzerland The legend on the right represents PCD cases per 100,000 inhabitants.

### Characteristics of the study population

Table 3 summarizes the basic characteristics of the 131 patients currently included in CH-PCD. Their median age is 25 years (range 5-73) and 51% are male. Four patients have died, one at the age of 45 from multi-organ failure following acute appendicitis and one at the age of 66 from acute respiratory failure. Date and cause of death were not registered in the charts for the other two patients. Diagnostic information was available for 92 patients, and age at diagnosis ranged from birth to age 54 years, with a median age of 4.6 years. Among these 92 patients, only 41 have a definite PCD diagnosis based on current international diagnostic guidelines [10]—primarily hallmark ultrastructural defects identified by electron microscopy (EM). Twenty-two patients were diagnosed only by low nasal NO (nNO) value and/or video microscopy analysis (VM), and the remaining 29 patients by clinical symptoms alone. Most of the patients in these two diagnostic groups have not undergone the full diagnostic algorithm proposed by international guidelines [10,18].

**Table 3.** Basic characteristics of PCD patients currently included in the Swiss PCD Registry (N=131)

|  | N (%) |
| --- | --- |
| **Sex** |  |
| Male | 67 (51) |
| Female | 64 (49) |
| **Vital status** |  |
| Alive | 119 (91) |
| Dead | 4 (3) |
| Lost to follow-up/moved abroad | 8 (6) |
| **Current age (N=127)<sup>¶</sup>** |  |
| 0-9 years | 18 (14) |
| 10-19 years | 33 (26) |
| 20-29 years | 28 (22) |
| 30-39 years | 14 (11) |
| 40-49 years | 12 (10) |
| ≥50 years | 22 (17) |
| **Laterality status** |  |
| Situs solitus | 50 (38) |
| Situs inversus | 37 (28) |
| Heterotaxia | 4 (3) |
| Information missing/not yet collected | 40 (31) |
| **Diagnostic information (N=92)** |  |
| Definite PCD diagnosis<sup>+</sup> | 41 (45) |
| Possible PCD diagnosis<sup>#</sup> | 22 (24) |
| Clinical diagnosis only | 29 (32) |
| **Age at diagnosis (N=92)<sup>&</sup>** | 4.6 (0-54) |

Data are presented as n (%) unless stated
<sup>¶</sup>As of May 2018, excluding the four deceased patients
<sup>+</sup>Defined as hallmark PCD electron microscopy findings and/or biallelic gene mutation identified based on the European Respiratory Society PCD Diagnostics Task Force guidelines
<sup>#</sup>Abnormal light or high-frequency video microscopy finding and/or low ( $\leq 77$ nL·min<sup>-1</sup>) nasal nitric oxide value
<sup>&</sup>In years, presented as median and range
Diagnostic information and age at diagnosis were available for 92 patients

### Current and ongoing studies with data from the Swiss PCD Registry

CH-PCD has participated in several international studies. It contributes Swiss data to the iPCD Cohort [13], which has been set up under the EU-funded FP7 Project BESTCILIA (Better Experimental Screening and Treatment for Primary Ciliary Dyskinesia, http://www.bestcilia.eu/). In addition to the 131 patients from Switzerland, the iPCD Cohort includes data from over 3700 PCD patients from 21 countries. CH-PCD patients have been included in a study of growth and nutrition in PCD patients from 16 countries since chronic respiratory disease can affect growth and nutrition, which can influence lung function [19]. The study found that PCD patients of all age groups are shorter than their peers and children younger than 10 years old, and also have lower body mass index (BMI). Results vary between countries, but height was reduced in most participating centres. Swiss patients had height and BMI within normal limits (Figure 4). Lung function was positively associated with height and BMI. Primary ciliary dyskinesia (PCD) has been considered to be a relatively mild disease, especially compared to cystic fibrosis (CF), thus another recent study has compared lung function in PCD and CF patients [20]. This study found that PCD patients from all countries, including Switzerland (Figure 4), had impaired lung function compared to normal references in all age groups. In children and adolescents, lung function of PCD patients was similar to that of cystic fibrosis patients (FEV1 % predicted was 81% in PCD patients versus 78% in CF patients aged 14-17 years). Lobectomy is suggested for the management of patients with localised bronchiectasis, so a third study (manuscript submitted) examines the effects of lobar resection in PCD across countries. The study shows that most lobectomised patients have poor lung function that continues to decline after surgery. In an effort to find ways to include more sensitive measures of lung function in routine clinical settings, another study aimed to assess alternatives to the lung clearance index (LCI) derived from a shorter protocol for the nitrogen multiple breath washout test (MBW) [21] . All patients included in CH-PCD with available MBW measurements were included with German patients in this study, which found a 79% prevalence of abnormal values for LCI2.5% and 72% for the significantly shorter LCI5%. Ongoing studies from the iPCD cohort (including CH-PCD data) describe the evolution of PCD diagnostic methods in Europe over the years and the neonatal manifestations in patients with PCD [22].

**Figure 4.**
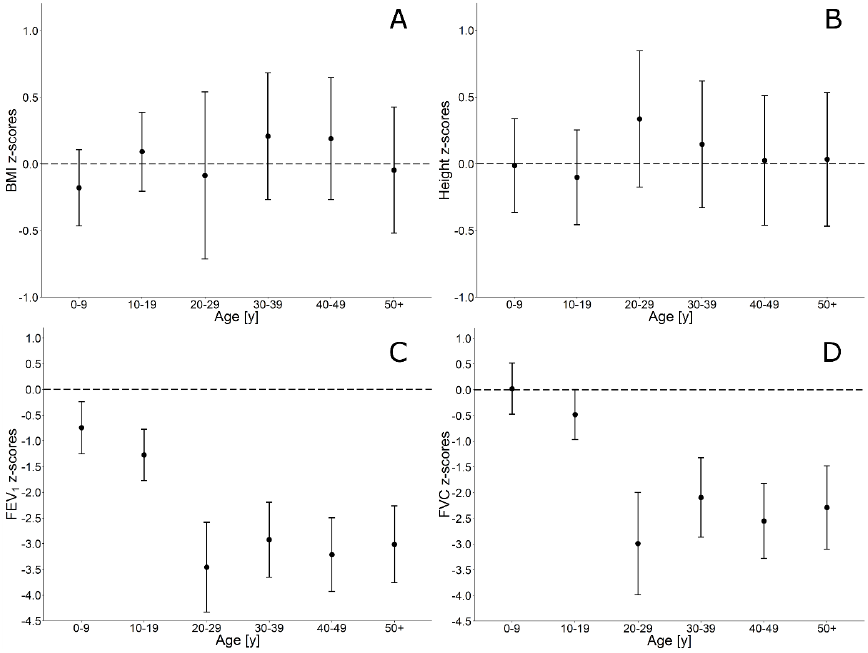
Growth and lung function of patients participating in the Swiss PCD registry Height (A) and BMI (B) by age group compared to national references; FEV1 (C) and FVC (D) by age group compared to GLI international references. All values are presented as mean z-scores (95% CI) after adjusting for sex and level of diagnostic certainty.

CH-PCD has also provided data to the International PCD Registry [12] and participated in the development of PCD-specific, health-related quality of life questionnaires (QoL-PCD). The QoL-PCD questionnaires are the first multidimensional measure to assess health-related quality of life. They were originally developed and validated in English in four different versions for children 6-12 years old, adolescents 13-17 years old, adults, and parent-proxies [23-25]. We translated them into German using words and phrases that are easily understood both in Germany and in Switzerland, and performed cognitive testing for the translated questionnaires in Swiss patients of different age groups. CH-PCD was also used to identify eligible patients for the first multicentre randomised controlled trial in PCD (trial registration: EudraCT 2013-004664-58) [26], which determined the efficacy and safety of azithromycin maintenance therapy. Outcome measures included number of exacerbations, lung function measured by spirometry, body plethysmography and multiple breath washout, QoL-PCD, hearing impairment, sputum microbiology and inflammatory markers. The study is now concluded and results will soon be available.

## Discussion

CH-PCD is one of few PCD patient registries worldwide and the only one that actively tries to identify all patients diagnosed in a single country (Switzerland). After contacting all medical specialties that could encounter PCD patients in the largest part of Switzerland, we found a) no awareness of PCD among practitioners who are not adult or paediatric pulmonologists, b) decentralized care and diagnosis of the disease, and c) low prevalence of diagnosed PCD patients, particularly among adults.

### Interpretation of results and comparison with other studies

Nearly all patients with PCD identified in Switzerland were reported by paediatric and adult pulmonologists whom we contacted. The low awareness of PCD among all other medical specialties is evident. This is understandable for nephrologists or neurologists, but is surprising for ENT specialists because ENT problems are as common—if not more so—as lower respiratory problems in PCD. In addition to low awareness, our data have confirmed that the management of PCD in Switzerland is decentralized and most physicians treat small numbers of patients.

The registry data confirm an earlier European study that found that children with PCD in Switzerland were treated in 17 centres (8 of which are tertiary) with 1 to 5 patients treated per centre [11]. In contrast to CF, which is treated in a few recognized centres, Switzerland has no recognized PCD centres, although the disease is even rarer, and many patients are managed by private practitioners. It is unclear how this affects the care of PCD patients in Switzerland. Even more importantly, PCD diagnosis is decentralised and Switzerland does not yet have designated diagnostic centres that perform all or at least most of the recommended diagnostic tests. This contrasts with other countries with one national PCD centre, such as Denmark, or a collaborative diagnostic network of centres as in the UK [27]. The lack of designated diagnostic centres could explain why fewer than half of the patients identified had a definite diagnosis based on international recommendations. The diagnosis of some patients were based only on clinical manifestations, while in others diagnosis was based on nNO measurement alone, which can have low positive predictive value, or on VM analysis on a single occasion instead of the recommended three samples, analysed in different occasions.

Reported prevalence of PCD varies importantly between studies. In a survey of the European Respiratory Taskforce (ERS TF) sent to all centres treating children with PCD, Switzerland had a prevalence of 1 in 20,000 among children aged 5-14 years, one of the highest among participating countries [28]. Our CH-PCD data suggest a prevalence of 1 in 27,000 for the age group of 5-14 years, but much lower prevalence of known cases for patients older than 30 years. The higher prevalence of disease diagnosed in adolescents aged 10-20 years compared to children aged 0-10 years and young adults aged 20-30 years suggests that most PCD patients are diagnosed late. We have made every effort to contact all of the physicians who might care for patients with PCD in most of Switzerland. Based on the low prevalence of disease diagnosed in adults older than 30 years, we conclude that a large proportion of adult patients with PCD are not yet diagnosed. Most older adults with PCD could have not been diagnosed as children when PCD was poorly understood and awareness of the disease was even lower. Our data suggest that these patients are also not diagnosed in adulthood; otherwise, the numbers of adult patients included in the registry should be much higher than the paediatric patients. This observation is in accordance with international data from the iPCD cohort and other available studies [13,29] suggesting that PCD was for many years considered a paediatric disease. Also, some adult pulmonologists believe that diagnosis of PCD would not change the management of their “non-CF bronchiectasis” patients, and they are therefore less diligent in following diagnostic guidelines. A third possibility for the small numbers of diagnosed adults could be increased mortality from PCD in adulthood. However, data on PCD mortality is lacking worldwide, and in our experience the severity of disease is highly variable but it is generally less severe in adults than cystic fibrosis [5,29,30].

### Strengths and limitations

The biggest strength of CH-PCD is the thorough methodology for patient identification, registration, and standardised data collection. We actively contact all physicians who might be involved in the management of patients with PCD using a uniform process. We also collaborate closely with a national network of clinical and diagnostic collaborators and advisors, and with the Swiss PCD patient organization. We present regular updates in national meetings and annual reports. The CH-PCD database allows data sharing and linking with international PCD projects such as the international PCD registry and the iPCD cohort (with high-level measures taken for data protection). The main limitation of the registry is that at time of patient registration, data collection is retrospective, which might lead to inconsistent and missing information for some patients. The CH-PCD team tries whenever possible to clarify these issues in communication with the treating physicians.

### Future implications for clinical management and research

The data from CH-PCD underline the current weaknesses of diagnosis and management of PCD in Switzerland. In the coming years, we need to increase the effort to raise the awareness of all physicians involved in the care of PCD, including nonrespiratory specialists. CH-PCD will continue to contact physicians nationwide from all specialties that are relevant to PCD not only to identify patients but also to inform them about the disease. CH-PCD activities and research results will be broadly presented in local and national meetings and conferences, and in both scientific and lay publications. At the same time, we need to catch up with the current international standards of PCD diagnosis and management and prioritize the establishment of centralized reference centres. Efforts are currently underway to establish a designated diagnostic centre at the University of Bern. Diagnostic approaches need to be updated and new patients should follow the available diagnostic algorithms [10]. Older patients should also benefit from recent diagnostic advances and should be invited to be retested whenever needed [29]. These efforts and advances should be communicated to the government and policymakers to allow health policies to be updated because, among other things, getting disability insurance for PCD in Switzerland is at present allowed only via diagnosis by EM analysis.

In an effort to improve prospective data collection, we developed adult and parent-proxy version versions of a standardised PCD follow-up form and a patient symptom questionnaire in collaboration with an international group of PCD experts including pulmonologists, paediatric pulmonologists, ENT specialists, diagnostic scientists, epidemiologists, study nurses, and physiotherapists.[16] The follow-up form includes key questions on diagnostic tests, signs and symptoms, clinical tests and treatments. It consists of several modules, including a more extensive module to be used at baseline and several shorter ones for regular follow-up visits. The follow-up form and the questionnaires are available in English, German, and French, and are currently being piloted in PCD clinics in countries using one of these languages.

### In summary

CH-PCD is now established in Switzerland and PCD patient identification is ongoing. This article highlights the value of the registry as a research tool in national and international collaborative studies and the need to improve clinical practice in Switzerland. Development of centralised diagnostic and management centres and adherence to international guidelines [10] are needed to improve diagnosis and management—particularly for adult PCD patients.

## Acknowledgements

We want to thank all the patients in the Swiss PCD registry and their families, and we are grateful to the Kartagener Syndrom und Primäre Ciliäre Dyskinesie e.V patient organisation, which closely collaborates with us. We thank all the physicians who helped identify patients and worked closely with us throughout building the Swiss PCD registry. We appreciate the work of Agatha Wisse and Anna Bettina Meier (Institute of Social and Preventive Medicine, University of Bern, Switzerland) and all the medical master’s students who contributed to patient identification and data collection : Sabrina Imsand, Petra Vogel, Marc Eich, Edwige Collaud, Anu Jose, Marion Amez-Droz, Lorenza Pacchin, Leonie Huesler and Michael Schoenholzer (Institute of Social and Preventive Medicine, University of Bern, Switzerland). We also thank Garyfallos Konstantinoudis (Institute of Social and Preventive Medicine, University of Bern, Switzerland) for his help with figure 3, and Christopher Ritter (Institute of Social and Preventive Medicine, University of Bern, Switzerland) for his editorial suggestions.

## **Current CH-PCD Working** Group (in alphabetical order)

Juerg Barben, St.Gallen; Jean-Louis Blouin, Geneva; Martin Brutsche, St. Gallen; Carmen Casaulta, Bern; Christian Clarenbach, Zurich; Regula Corbelli, Geneva; Lin Dagmar, Bern; Reta Fischer, Bern; Thomas Geiser, Bern; Myrofora Goutaki, Bern; Florian Halbeisen, Bern; Juerg Hammer, Basel; Andreas Jung, Zurich; Claudia Kuehni, Bern; Philipp Latzin, Bern; Romain Lazor, Lausanne; Marco Lura, Luzern; Alexander Moeller, Zurich; Carlo Mordashini, Bern; Jean-Claude Pache, Lausanne; Nicolas Regamey, Luzern; Bernhard Rindlisbacher, for the Swiss PCD patients organization; Isabelle Rochat, Lausanne; Stefan Tschanz, Bern; Agatha Wisse, Bern; Maura Zanolari, Bellinzona.

